# Rapid decline of prenatal maternal effects with age is independent of postnatal environment in a precocial bird

**DOI:** 10.1101/2023.05.26.542383

**Authors:** Oscar Vedder, Barbara Tschirren, Erik Postma, Maria Moiron

**Author notes:** **Author contributions** Conceptualization and experimental design: OV, BT and MM. Data collection: OV. Data analysis: MM, with feedback from OV, EP and BT. Writing: OV and MM, with feedback from EP and BT.

## Abstract

Maternal effects are an important source of phenotypic variation with potentially large fitness consequences, but how their importance varies with the quality of the environment across an individual’s ontogeny is poorly understood. We bred Japanese quail (*Coturnix japonica*) of known pedigree and experimentally manipulated the quality of offspring diet, to estimate the importance of prenatal maternal effects in shaping variation in body mass from hatching to adulthood. Maternal genetic effects on body mass at hatching were strong, and largely caused by variation in egg mass, but their importance rapidly declined with age. Whereas there was a large effect of diet on growth, this did not affect the decline of maternal effects variance. The importance of additive genetic and residual variance increased with age, with the latter being considerably larger in the poor diet treatment. Hence, we found no evidence for prenatal maternal effect by postnatal environment interactions, and that prenatal maternal effects are rapidly replaced by direct additive genetic and residual effects when offspring start to develop outside the egg. Thereby these results shed new light on the dynamics of the role of maternal versus offspring genes across ontogeny and environments.

## Introduction

Mothers do not only provide genetic material to their offspring, but also determine a large part of their offspring’s developmental environment. The phenotype of an individual can therefore be affected by the phenotype of the mother, independent of the effect of the genes she passed on to her offspring (Mousseau and Fox 1998, Wolf and Wade 2009). Such effects are generally regarded as ‘maternal effects’ and can be quantified as the effect of maternal traits on offspring trait values (trait-based approach), or as the proportion of total phenotypic variance among offspring that is explained by maternal identity, over and above the effect of their shared genes (variance-component approach) (Falconer and Mackay 1996, McAdam et al 2014).

Maternal effects are potentially of great importance in organismal biology, as they can have lasting effects on individuals throughout life (Moore et al. 2019). They may mediate a phenotypically plastic response to environmental change, with mothers adjusting their offspring phenotype to the future environment (Marshall and Uller 2007, Kuijper and Hoyle 2015). Furthermore, they can affect the speed of evolutionary processes, through positive or negative correlations between offspring and maternal genetic effects (Kirkpatrick and Lande 1989, Wolf et al. 1998, McAdam et al. 2002, Räsänen and Kruuk 2007, McGlothlin and Galloway 2014, Pick et al. 2019).

The relative importance of maternal effects versus offspring genes inherited from their mother can be difficult to estimate. For example, large mothers may pass on genes for large size, but their size may also allow them to provide their offspring with a more favourable developmental environment (Sinervo 1993, Krist and Remeš 2004). While for maternal effects that are caused postnatally, both effects can be uncoupled by exchanging offspring between mothers (cross-fostering), maternal effects may already act during embryonic development, with organizational processes in the embryo affected by prenatal maternal provisioning of resources and developmental cues (Henry and Ulijaszek 1996). At this developmental stage, cross-fostering is not feasible, and prenatal maternal effects need to be separated from offspring genetic effects in other ways.

Phenotypic data on individuals of known relatedness allows, at least in theory, for statistically disentangling the amount of variance in an offspring trait that can be attributed to maternal effects (environmental and genetic) versus direct offspring genetic effects, using quantitative genetic mixed (or ‘animal’) models (Wilson et al. 2005, Kruuk and Hadfield 2007). In practice, the power of this approach is constrained by the amount and structure of the data. In this respect, the presence of paternal half siblings (same father, different mother) greatly facilitates the estimation of maternal effects. Yet, in natural populations of some taxa, such cases may be too rare, or systematically biased with regard to offspring genotype or environment, for the reliable estimation of maternal effects (Kruuk and Hadfield 2007). In such cases, controlled breeding in captivity allows for the creation of a pedigree structure that does not suffer from such shortcomings (e.g. Pick et al. 2016a).

The downside of captive breeding experiments is a loss of ecological realism. For example, captive individuals are often raised in a favourable and highly standardized environment. To what extent the environment modulates the strength of maternal effects remains to be evaluated, but maternal effect by environment interactions may be common (Charmantier and Garant 2005, Rowinski and Rogell 2017). For example, a maternal effect on pup growth rate in North American red squirrels (*Tamiasciurus hudsonicus*) was significantly reduced when extra food was provided (McAdam and Boutin 2003). In the soil mite (*Sancassania berlesei*), a negative effect of maternal age on offspring reproductive performance was stronger at lower densities (Plaistow and Benton 2009). Finally, in brown trout (*Salmo trutta*), a positive effect of egg size on offspring performance was found to be larger in poor environments (Einum and Fleming 1999). Only a few studies, however, have specifically tested for maternal-effect-by-environment interactions using an animal model that captures the resemblance among offspring from the same mother over and above the resemblance caused by direct additive genetic effects, combined with controlled breeding and experimental manipulation of the environment during offspring growth (Vega-Trejo et al. 2018, Vrtílek et al. 2021). In a captive population of mosquitofish (*Gambusia holbrooki*) significant maternal-effect-by-environment interactions were found, with stronger maternal effects on female age at maturity in a low food environment and on male size at maturity in a standard food environment (Vega-Trejo et al. 2018). On the other hand, in seed beetles (*Callosobruchus maculatus*) maternal effects on several life-history traits did not differ between rearing environments, and maternal effects were negligible in both the original and a novel environment (Vrtílek et al. 2021).

In this study we test if the importance of prenatal maternal effects on offspring body mass during growth varies with the quality of the early-life environment in a precocial bird, the Japanese quail (*Coturnix japonica*). To this end, we made reciprocal crosses between lines originally selected for low and high maternal investment in egg size (Pick et al. 2016b), creating full- and half-siblings of which the exact ancestry was known for >10 generations. With this breeding design we built on previous work by Pick et al. (2016a), who found maternal effects to decline, but not disappear, across the first four weeks of their life. By this age, they are independent of parental care, but not yet fully grown and sexually mature. In particular, we raised hatchlings on either a poor or a standard rearing diet, and to test if maternal effects on body mass during growth persist in adult life, we monitored body mass over the complete growth phase (until 12 weeks old). As such, the primary objective of the study was to explore the evolutionary genetics of maternal effects on offspring body mass in different early-life environments. Specifically, we estimated how much variation in offspring mass is due to maternal effect and additive genetic variance, and how this depends on age and diet. In a second step, we quantified the role of egg mas in causing this variation.

## Methods

### General procedures and data collection

We used lines of domesticated Japanese quail divergently selected for ‘high’ and ‘low’ maternal investment in egg mass (for details see Pick et al. 2016b). Both selection regimes where replicated, such that there were four lines in total. These lines were originally established at the University of Zurich, Switzerland, but transferred to the Institute of Avian Research in Wilhelmshaven, Germany, in 2017. There they were kept in large outdoor aviaries, and selectively bred in pairs once a year to maintain the lines. This has resulted in a pedigree that for the purpose of this study encompassed 12 generations (see details below).

In 2019 and 2020, 1-year-old males and females of each line were both mated twice; once with a partner of the same line and once with a partner of the alternate selection lines, resulting in both maternal and paternal half-siblings. Mating took place in breeding cages (122 cm × 50 cm × 50 cm), in which pairs spent 10-14 days together. One-year-old female quail lay eggs at a rate of about 0.8 eggs per day (Vedder et al. 2022), and all eggs were collected daily. Upon collection, eggs were marked and weighed to the nearest 0.01 g. The first eggs to be fertilized by the paired male were laid two days after pairing. At least 14 days prior to pairing, females were housed in single sex aviaries to prevent stored sperm of previous males to fertilize any eggs (sperm storage in Japanese quail is max. 10 days; Birkhead and Fletcher 1994). The order of mating (same line vs. alternate line) was randomized. Eggs were stored at 12° C, and artificially incubated in batches within 14 days after laying.

Incubation was performed at 37.7° C and 50% relative humidity in fully automatic incubators (Grumbach, ProCon automatic systems GmbH & Co. KG, Mücke, Germany) that turned eggs every hour for the first 14 days of incubation. After 14 days of incubation, eggs were placed in marked individual compartments to allow the hatchling to be linked to the egg it hatched from. Further incubation was done at 37.2° C and 70% relative humidity, without egg turning, in hatching incubators (Favorit, HEKA Brutgeräte, Rietberg, Germany). From 16 days of incubation onwards, the hatching incubators were checked once a day for new hatchlings until no viable eggs were left (around 19 days after incubation).

At hatching, chicks were marked with a numbered plastic leg ring. They were weighed to the nearest 0.01 g and placed in heated rearing cages (109 × 57 × 25 cm, Kükenaufzuchtbox Nr 4002/C, HEKA Brutgeräte, Rietberg, Germany), with a maximum of 30 individuals per cage. Hatchlings were randomly distributed over rearing cages with one of two rearing diets differing in quality, in such a way that offspring from the same parent combination were roughly equally divided over the two treatments. What is referred to from now on as the poor-quality diet contained 14.5% protein, 4.0% fat and 1.0% calcium, amongst others (see Online Supplementary Material for a complete overview), and had a caloric value of 11.4 MJ/Kg. What we refer to as the standard diet contained 21.0% protein, 4.0% fat and 1.1% calcium, amongst others (see Online Supplementary Material for a complete overview), and also had a caloric value of 11.4 MJ/Kg. Both are commercially available poultry feeds (GoldDott, DERBY Spezialfutter GmbH, Münster, Germany). The lower protein content of the poor-quality was chosen to substantially reduce growth rate, without compromising chick survival (Weber and Reid 1967). The protein content of the standard diet was chosen to have a high growth rate, without leading to impairments associated with too rapid growth (Weber and Reid 1967). All chicks received *ad libitum* food and water, and the food was ground to a homogeneous powder to prevent selective eating and to ensure the consistency of both diets was similar. The temperature of all rearing cages was set at 37.0° C at hatching and gradually lowered to 20-25° C over the course of three weeks. All chicks were kept at a 16-8 h light-dark cycle. The plastic leg rings were replaced with uniquely numbered aluminum rings when chicks were between 14 and 35 days old, depending on the size of their legs. After three weeks in the rearing cages, the chicks were transferred to outdoor aviaries equipped with heat lamps, where they were kept on the same rearing diets as they were on before, and received a minimum of 16 h of daylight.

All chicks were weighed at weekly intervals after hatching (= age 0 days) to the nearest 0.01 g at 7 days, and to the nearest g at all subsequent ages until 84 days old. Chicks were sexed based on plumage characteristics, which are distinctly different between males and females after about 4 weeks old. From 5 weeks old onwards, birds were individually checked for the onset of reproductive activity every 2-3 days. For males this was done by checking for the production of cloacal foam (Sachs 1969). For females this was done by checking for the presence of an egg in the oviduct through palpation. The average age (± SD) of onset of reproductive activity was 44.7 (± 8.4) days and 51.5 (± 9.8) days for males and females with the standard diet, respectively, and 68.7 (± 14.5), and 74.1 (± 12.1) days for males and females with the poor diet, respectively. Sexually mature individuals from both treatment groups were housed together and provided with an adult layer diet, with 19.0% protein, 4.6% fat and 4.8% calcium, amongst others, and a caloric value of 9.8 MJ/Kg (GoldDott, DERBY Spezialfutter GmbH, Münster, Germany).

Chick mortality was low (5.7%), mostly occurred in the first few days after hatching, and did not significantly differ between the diet treatments (poor diet: 6.8%, standard diet: 4.6%; Table S1). Yet, to prevent changes in maternal effects over ontogeny due to selective disappearance rather than within-individual changes over ontogeny, we excluded all chicks that died before 84 d old from analyses (*n* = 39). This left a dataset of 648 (353 in 2019, 295 in 2020) individuals with complete growth trajectories, that were produced by 144 (82 in 2019, 62 in 2020) unique parent combinations, with 86 unique fathers and 78 unique mothers.

### Pedigree

The pedigree traced the ancestry of all offspring back to the unselected founder (or base) population and had a maximum depth of twelve generations. We pruned the pedigree to individuals with measured phenotypes and all their known ancestors using the R package *nadiv* (v. 2.17.1, Wolak 2012). In total, the pruned pedigree comprised 1414 records (data observations), and 1239 paternities and maternities. The numbers of full, maternal and paternal sibling pairs were 1969, 3152 and 3515, respectively. We calculated the inbreeding coefficient of all individuals with the R package *nadiv* (v. 2.17.1, Wolak 2012).

### Statistical analyses

We fitted a series of animal models to test whether prenatal maternal effects on offspring body mass vary with the quality of their early-life diet. First, we tested for the phenotypic effect of rearing diet on body mass across all age classes while accounting for the relatedness among individuals. To do so, we fitted a univariate animal model with body mass as response variable modelled with a Gaussian error distribution. We included random intercepts for individual, pair, mother and father identities. We also fitted a random intercept effect for individual identity linked to the relatedness matrix (additive genetic variance). As fixed effects, we fitted age (categorical variable with 13 levels, ranging from 0 to 84 days at a weekly interval), diet (categorical variable with two levels: standard and poor), sex (categorical variable with two levels: male and female), year (categorical variable with two levels: 2019 and 2020), individual inbreeding coefficient (continuous variable), paternal and maternal selection line (categorical variables with two levels: ‘high’ and ‘low’) and their interaction, maternal and paternal line replicate (categorical variable with two levels: 1 and 2), and the 2-way interactions of age with sex and diet, and the 3-way interaction between mother and father line and diet.

Second, we tested for an effect of early-life diet on maternal (genetic and environmental) effects for each age class. To do so, we fitted 13 bivariate animal models with body mass at a specific age (age ranged from zero to 84 days with weekly intervals) for each diet treatment as response variables. These age-specific body masses were mean-centered and variance-standardized within their diet treatment and modelled with a Gaussian error distribution. The bivariate animal models included random intercepts for individual and maternal identity linked to the relatedness matrix (additive genetic variance, V_A_ and maternal genetic variance V_MATERNAL GENETIC,_ respectively). We also fitted mother and father identity not linked to the pedigree to account for repeated measures across mothers and fathers, and to estimate the maternal and paternal permanent effects (V_MATERNAL ENVIRONMENTAL_ and V_PATERNAL ENVIRONMENTAL_, respectively), capturing the resemblance among offspring from the same mother and father over and above the resemblance caused by direct additive genetic and indirect maternal genetic effects. Additionally, we modelled the covariance between treatments across all the random effect parameters (i.e. additive genetic covariance, COV_A_; maternal genetic covariance, COV_DAM_; maternal permanent covariance, COV_MOTHER_; and paternal permanent covariance, COV_FATHER_. The residual covariance across treatments is inestimable because individuals are only ever present in one diet group. As fixed effects we fitted, for both response variables separately across all models, sex (categorical variable with two levels: male and female), year (categorical variable with two levels: 2019 and 2020), and individual inbreeding coefficient (continuous variable). To specifically estimate both age-specific additive genetic effects and parental effects for the unselected ‘founder’ population, we corrected for phenotypic differences in age-specific body mass between the lines and their replicates that may have occurred as a correlated response to artificial selection on egg mass (Pick et al. 2016b). To this end we added maternal selection line (categorical variable with two levels: ‘high’ and ‘low’), paternal selection line (categorical variable with two levels: ‘high’ and ‘low’), the interactive effect of maternal and paternal line, mother line replicate (categorical variable with two levels: 1 and 2), and father line replicate (categorical variable with two levels: 1 and 2) as fixed effects.

Third, after we identified and quantified at which ages there was evidence of maternal genetic effects (i.e. week 1 and week 2), we built a hybrid model that applies both a variance-partitioning and a trait-based approach to maternal effects (McAdam et al 2014, Pick et al 2016a). This model allowed for distinguishing between maternal genetic effects mediated by maternal egg investment and other forms of maternal inheritance (e.g. due to a maternal genetic correlation between egg size and offspring traits via maternally inherited genes such as mitochondrial genes). This model tested for the effects of fresh egg mass, partitioned in line, line replicate, average mother and within mother effects, on offspring body mass. We fitted bivariate animal models where the response variables were individual body mass at the two diet treatments at week 1 or 2. Age-specific body mass was mean-centered and variance-standardized within the diet treatment and modelled with a Gaussian error distribution. We modeled the same random effect structure as in the animal models described above. As fixed effects, we again fitted sex, year and individual inbreeding coefficient. However, instead of correcting for selection line effects, we directly quantified the effect of the fresh mass of the egg an individual hatched from. We did this with a within-subject centering approach (van de Pol and Wright 2009), by including the average fresh egg mass for both maternal lines averaged across replicates (‘average line egg mass’ for ‘high’ and ‘low’), the deviation from these two means for each replicate (‘delta replicate’ for ‘high1’, ‘high2’, ‘low1’ and ‘low2’), the deviation from these four replicate mean values for each individual mother (‘delta mother’ for 78 unique mothers), and the deviation from these 78 values for each individual egg (‘delta within mother’, 648 unique eggs) as fixed effects. This way, the fresh mass of each egg can be calculated as ‘average line egg mass’ + ‘delta replicate’ + ‘delta mother’ + ‘delta within mother’, yet all these underlying variables are uncorrelated and should thus have the same effect on offspring body mass when fitted together in one model, if the effect of fresh egg mass is causal (see Pick et al. 2016a for a similar approach).

All models were fitted using a Bayesian framework implemented in the statistical software R (v. 4.1.1, R Core Team 2021) using the R package *MCMCglmm* (Hadfield 2010). For all models we used parameter-expanded priors (Hadfield 2010). The number of iterations and thinning interval were chosen for each model so as to ensure that the minimum MCMC effective sample size for all parameters was 1000. Burn-in was set to a minimum of 2000 iterations. The retained effective sample sizes yielded absolute autocorrelation values lower than 0.1 and satisfied convergence criteria based on the Heidelberger and Welch convergence diagnostic (Heidelberger and Welch 1981). We drew inferences from the posterior mode and 95% credible intervals (95% CI). Variance-standardized components, such as heritabilities (h^2^), were conditional on the variance explained by fixed effects and estimated as the proportion of the total phenotypic variance explained by the focal variance component. Mean-standardized components, such as evolvabilities (I_A_), were estimated as the focal variance component divided by the relevant squared mean body mass (Houle 1992; Hansen et al. 2011).

## Results

### Effect of diet treatment on body mass

There was a large effect of diet on mean body mass across age classes. At all ages after hatching, the poor-quality diet led to considerably lighter average body mass, which lasted until all individuals were full grown (Fig. 1, Table S2). Offspring body mass was also dependent on sex, as after 28 days females became heavier than males (Fig. 1, Table S2). In terms of random effect components, we observed that individual identity (including both additive genetic and permanent environmental effects) explained most of the phenotypic variation in offspring body mass (∼57%), indicating that individuals were repeatable in their body mass across ages (Table S2).

**Figure 1.**
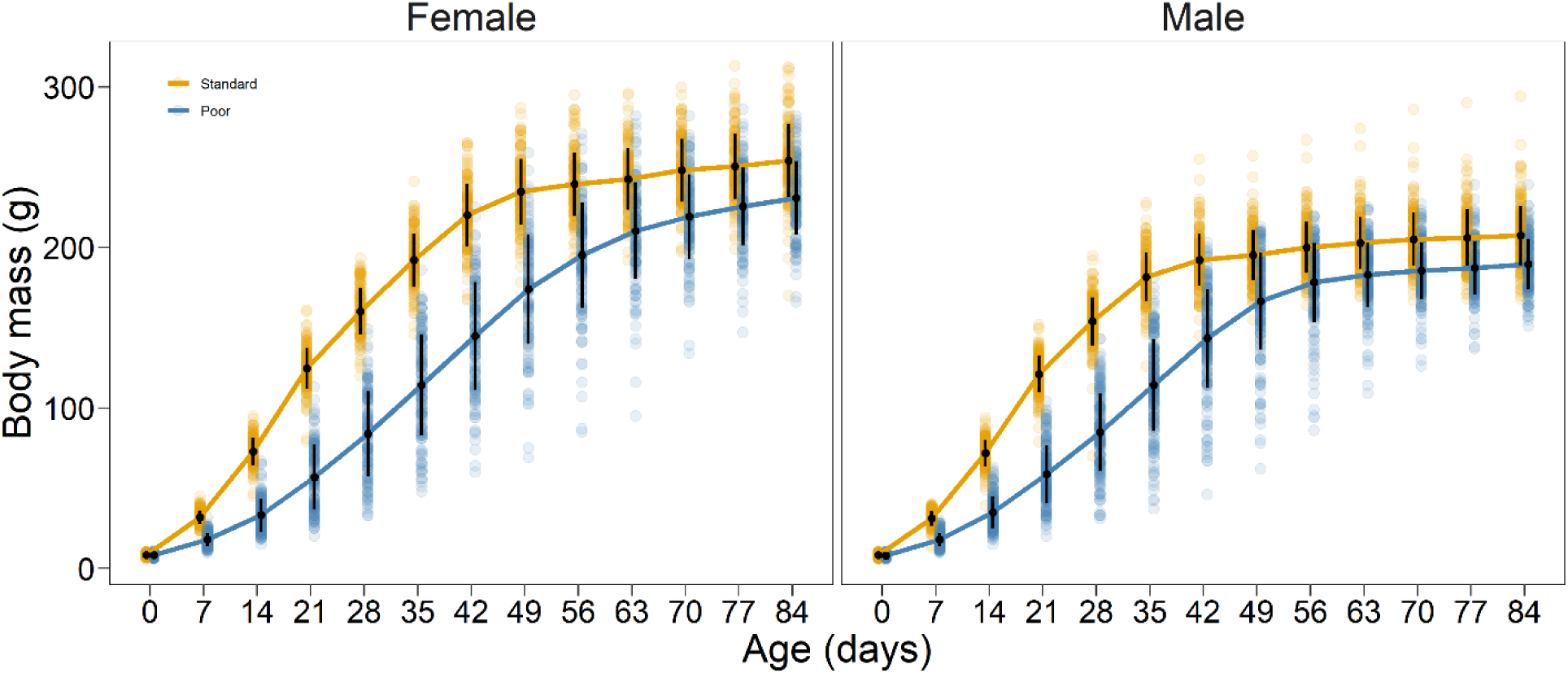
Sex- and rearing diet-specific postnatal growth in body mass of Japanese quail chicks. Black dots with error bars represent age-specific means ± standard deviations, and the coloured dots represent all individual datapoints. The difference between rearing diets is significant for all ages after zero (see Table S2).

### Test for diet-specific maternal effects on body mass

The maternal genetic variance was relatively large at hatching, representing ca. 61% of the total phenotypic variance in body mass in both diet treatments (Fig. 2, Table S3). At this point, the maternal genetic effects were also strongly correlated across the two diet treatments (r ∼1, Table S3), suggesting that the same maternal genes cause the maternal effects in both treatment groups. This is as expected, because at hatching chicks were weighed before receiving food, and therefore were not yet exposed to different environments. However, in the following age classes, the maternal genetic effect decreased to become effectively zero (Fig. 2, Table S3), making it impossible to reliably estimate the maternal genetic (co)variances within and across treatments (Table S3). Regardless, we did not observe any clear differences in effect sizes of maternal genetic effects between the two diet treatments at any age class (neither in absolute, variance-standardized, nor mean-standardized estimates, Fig. 2, Table S3).

**Figure 2.**
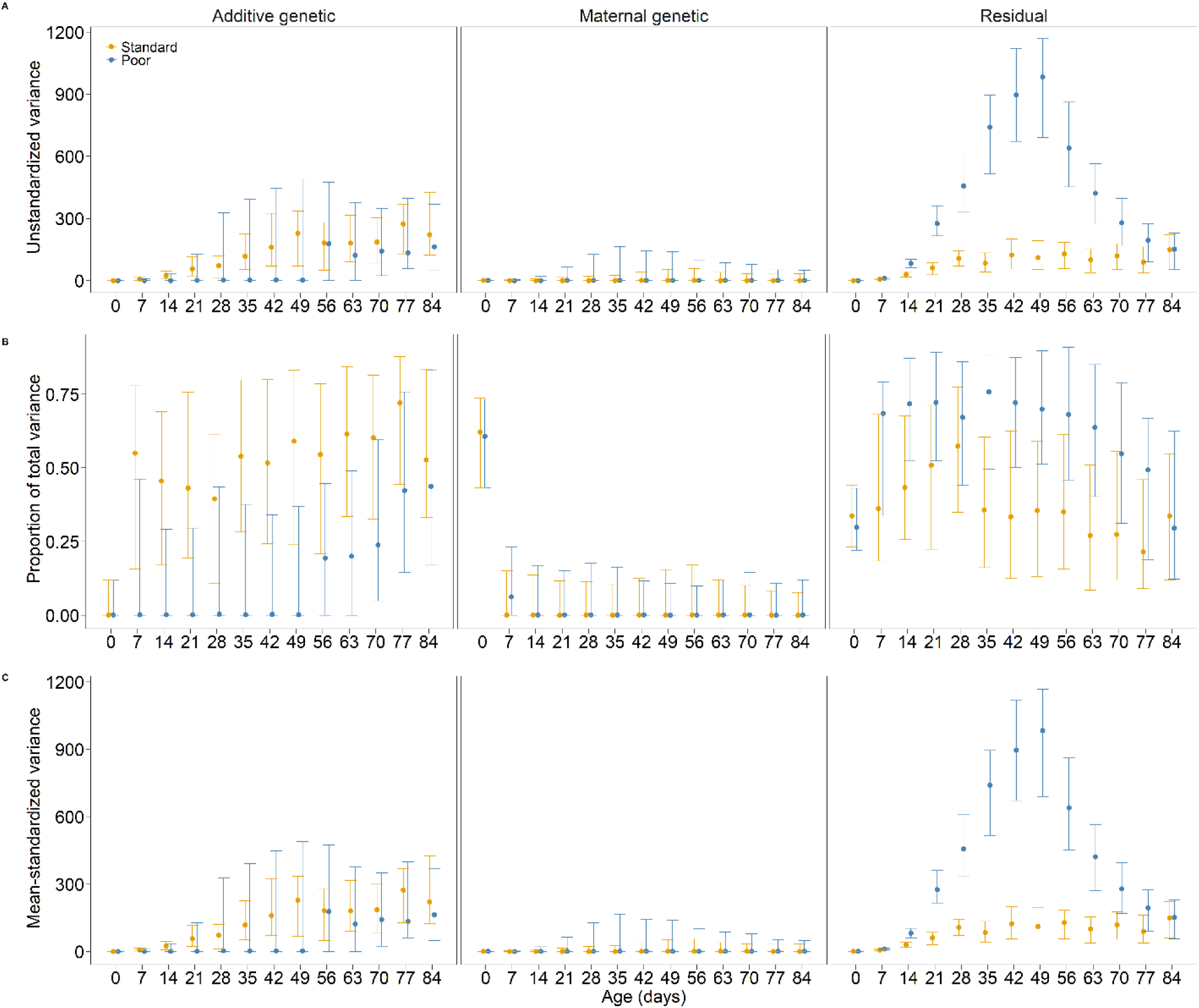
Age-specific additive genetic variance, maternal genetic variance, and residual variance in body mass of Japanese quail chicks (from left to right), presented as A) unstandardized variance, B) proportional to the total variance (h^2^, m^2^, and r^2^), and C) mean-standardized variance (i.e. I_A_), subdivided over the two diet treatments. Dots with error bars represent the mean estimates with 95% credible intervals (see also Table S3).

Maternal and paternal permanent (environment) effects across all ages were not significantly different from, and close to, zero, and did not differ across diet treatments (Table S3). Additive genetic variance, heritability and evolvability of body mass increased with age and were larger in the standard diet treatment, although there were no consistently strong differences between treatments (i.e. the differences between diets were significant at days 21 and 63, but not for any other age class, Fig. 2, Table S3). However, residual variance in body mass was highest at intermediate age classes (roughly between days 35 and 56), especially in the poor diet treatment (Fig. 2, Table S3).

In terms of fixed effects, there was an important effect of the maternal line in all age classes, with offspring from high-investment mothers having, on average, heavier body mass (Table S3). There was no clear effect of paternal line at the early age classes (day 0 and 7), but it increased at later age classes (day 14 onwards), with it being significant but not clearly different from the maternal line effect (Table S3). The similar effects of the parental lines can be explained by genetic differences in body mass between the lines, with the early effect of maternal line caused by mothers from the high investment line also laying larger eggs (see below). The interaction between maternal and paternal line was not significant at any of the age classes, except at day 21 (Table S3), indicating that there is little evidence for non-additive effects of the parental lines. In none of the age-specific models was there a clear effect of year of experiment, and mother or father line replicate (Table S3). The individual inbreeding coefficient only had a negative effect on body mass at intermediate ages (from 14 to 35 days, Table S3), suggesting that inbreeding depresses the rate of growth, but not the initial and final body mass.

### Hybrid model to identify the underlying mechanism of maternal effects

The effects of fresh egg mass, partitioned in line, line replicate, average mother and within mother effects, on body mass at hatching were strong, and not statistically different from each other (Table S4), which is in line with the relationship between egg mass and offspring mass being causal. By including these components of egg mass variation as covariates in the animal model (Table S3), we observed a strong decrease in the variance explained by maternal genetic effects at hatching (Fig. 3, Table S4). This decrease was strong to the extent that maternal genetic effects were not significantly different from zero anymore. As such, the addition of those sources of variation in egg mass explained all the important maternal genetic variance in body mass, pointing again towards a causal link. The same ‘hybrid’ model was also applied to body mass at 7 days, but here the sources of variation in egg mass tended to have different effects (Table S4). The particularly strong ‘delta replicate’ effect may be explained by the line replicates that laid large eggs for their line also being genetically larger in body mass at this age. Although there were no significant maternal genetic effects at this age, the correction for the sources of variation in egg mass further reduced any indication of maternal genetic variance in body mass (Fig. 3).

**Figure 3.**
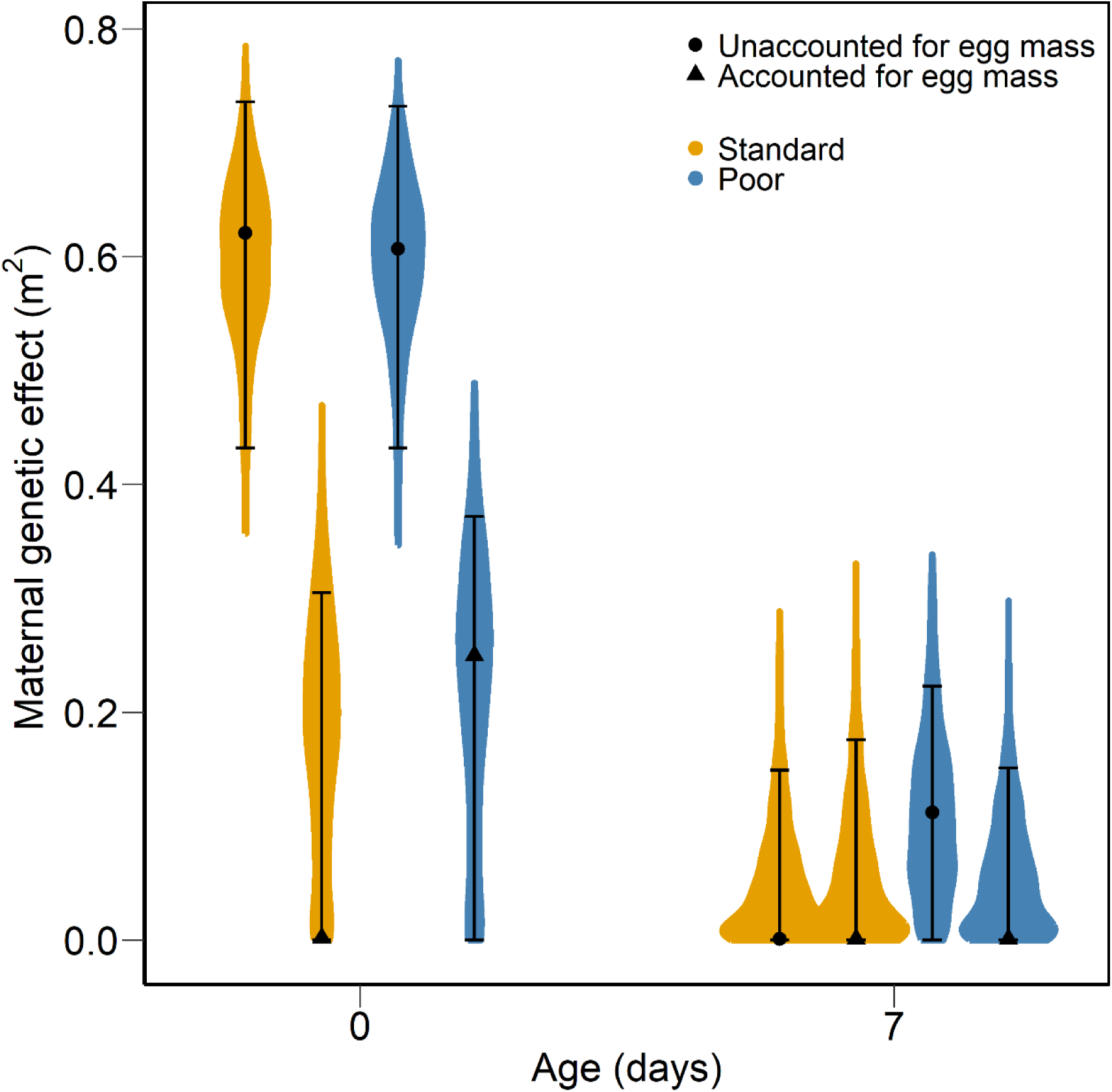
Estimates of rearing diet-specific maternal genetic effects (m^2^) on body mass at the ages of 0 and 7 days, before and after accounting for all sources of variation in the fresh mass of the eggs from which the chicks hatched. Symbols with error bars represent the mean estimates with 95% credible intervals (see also Table S3 and S4), and the coloured backgrounds the posterior distributions of the estimates.

## Discussion

Maternal effects can act as an important source of phenotypic variance and have evolutionary consequences, but to what extent this is environment-dependent remains poorly understood. Here we used controlled breeding and an experimental manipulation of the offspring diet during growth to explicitly test for prenatal maternal effect by environment interactions on body mass over ontogeny in Japanese quail. Although this approach, combined with quantitative genetic animal model analyses, is very useful in quantifying the true strength, evolutionary consequences, and environmental dependency of maternal effects, it has been rarely applied (Vega-Trejo et al. 2018, Vrtílek et al. 2021), and, to our knowledge, never before in birds.

We found strong maternal genetic effects on the body mass of a chick at hatching, yet this was before the chicks were exposed to the food treatment. Maternal environmental effects did not explain variation in body mass at any age class, or under any of the two diets. After the chicks started to grow, the maternal genetic effects on body mass quickly diminished under both rearing diets, despite growth being severely depressed by the poor-quality diet. There was, however, much more phenotypic variance in body mass under the poor-quality diet, which was not due to an increase in the variance attributable to parental or additive genetic effects.

Independent of diet, the strength of maternal effects on body mass declined faster than in a previous study using the same breeding design and selection lines of Japanese quail (Pick et al. 2016a). This difference can be explained by the fact that in the current study we controlled for ‘fixed’ differences between the lines in the models. This way we corrected for differences in selection on egg mass between the lines, and thereby estimated the strength of maternal effects in the ‘founder’, unselected, population. Hence, without artificially exaggerated genetic variance in egg mass, maternal effects were weaker and disappeared faster with age. There was, however, also variance in egg mass among, and within, mothers within the replicated lines, and this variance could almost completely account for the maternal effects on body mass at the earliest chick ages. These results are very similar to a study on blue tits (*Cyanistes caeruleus*) that found prenatal effects on post-hatching body mass to be almost fully explained by variance in egg mass (Hadfield et al. 2013). Hence, variation in the mass of eggs from which the chicks hatch is likely to be the most important cause of prenatal maternal effects on early-life body mass. Only traits that would be highly correlated with individual egg mass, similarly among and within mothers, could alternatively account for the prenatal maternal effects on body mass observed in our study.

The decline in maternal effects with age is in line with maternal effects generally explaining less variance in adult traits than in juvenile traits (reviewed by Moore et al. 2019). As such, the importance of the developmental environment that the mother provides most likely wanes with age. However, in laboratory studies such as ours, food is typically provided *ad libitum*, causing little competition over food. In the wild, where food is limited, it is possible that an early-life competitive advantage (e.g. larger size) caused by maternal effects, could lead to a positive feedback loop in resource acquisition and thereby also determine late-life performance (Fokkema et al. 2021). In addition, in the wild mothers may also determine post-hatching resource acquisition (via choice of oviposition site or by direct food provisioning) potentially causing a greater importance of postnatal maternal effects. Indeed, although we did not treat the post-hatching diet as a maternal effect, the chicks that received the poor diet remained of lower body mass at adult age (Fig. 1), which may have life-long performance consequences. A study on free-living red deer (*Cervus elaphus*), however, only found maternal effects on neonatal traits and not on adult traits (Gauzere et al. 2020), suggesting that also in the wild the importance of maternal effects may decrease over ontogeny.

Studies that synthesized estimates of maternal effects in relation to environmental quality tentatively suggest that the strength of maternal effects does not differ systematically between poor- and good-quality environments (Charmantier and Garant 2005, Rowinski and Rogell 2017). In this study we specifically conducted an experiment to test if the strength of prenatal maternal effects differed between different quality postnatal environments. Despite the chicks with the poor-quality diet weighing, on average, less than halve of those with the standard diet during peak growth, the strength of maternal effects did not decline noticeably slower in the poor environment. We thus did not find evidence for maternal (genetic or environmental) effects by environment interactions on growth. Since the strength of maternal effects is relative to the total variance in a trait, and trait variance may increase with trait mean, we also estimated the age-specific maternal effect variance divided by the squared average age-specific body mass (conceptually similar to evolvability; Houle 1992). If the absolute amount of maternal effect variance does not change with mean body mass, but other sources of variance (additive genetic and residual) do increase in concert with mean body mass, this should lead to higher estimates of this measure (Maternal genetic I_A_) in the poor diet environment. However, this was not the case, as with both diets these estimates were negligible already at seven days of age (Fig. 2). Instead, although the absolute amount of variance in body mass caused by maternal effects did not increase after hatching with both diets (Fig. 2), the residual variance increased rapidly with age with the poor diet, while the increase in additive genetic variance was slower with age, compared to the standard diet.

The extreme increase in residual variance in body mass with the poor diet also considerably lowered the proportion of the variance in body mass attributable to additive genetic variance, i.e. the heritability (V_A_/V_P_) during growth. This was unexpected, as despite being a topic of great interest, there appears to be no general pattern of how heritability depends on environmental quality across studies (Charmantier and Garant 2005, Rowinski and Rogell 2017). While the poor diet clearly exaggerated among individual differences in growth rate, it is difficult to identify the source of this residual variance. Such large differences in growth rate may be expected if there is intense competition for food, causing large disparity between ‘winners’ and ‘losers’. However, food was provided *ad libitum* and no obvious competition over food was observed. Moreover, as a larger initial body size would be likely to provide an advantage in competition over food, this should have increased the strength of maternal effects, and not of residual effects, because body mass at hatching was predominantly caused by maternal effects. Instead, we speculate that these large differences in growth rate may be caused by among-individual variation in protein synthesis efficiency (Marks 1993). Perhaps such variance is caused by non-additive genetic effects, potentially causing the animal model to allocate it to the residual variance component (Kruuk 2004, Class and Brommer 2020). Alternatively, for some reason the poor diet may have exaggerated stochastic effects on growth, or there were very subtle, unknown, environmental differences with a large effect on growth, particularly within this diet treatment.

In conclusion, our experimental test for prenatal maternal effect by environment interactions on body mass over ontogeny found no evidence for maternal effect by environment interactions at any age. There was only a strong prenatal maternal effect on mass at hatching, most likely caused by variation in egg mass, but this effect quickly diminished with age. We suggest that as soon as chicks start to grow, the among individual variance in body mass is primarily explained by additive genetic and residual effects (non-additive genetic, environmental or stochastic), of which the relative importance can vary considerably with the quality of the rearing diet.

## Acknowledgements

We thank Nils Becker and Kyra Fastenau for help with data collection, and Adolf Völk for the quail husbandry. The study was funded by grant number 428800869 from the German Research Foundation (DFG, Deutsche Forschungsgemeinschaft) to OV. MM was funded by an Alexander von Humboldt Research Fellowship for Postdoctoral Researchers. All practical procedures complied with current German law.

## Data Availability

The data will be deposited in Dryad upon acceptance of the manuscript.

