## Supplementary Material for "Rapid decline of prenatal maternal effects with age is independent of postnatal environment in a precocial bird"

**—Online Supplementary Material—**

### Detailed description of the two rearing diets

**Standard diet:**

Ingredients: Crude protein 21.0%, crude fat 4.0%, crude fibre 3.5%, crude ash 7.0%, lysine 1.1%, methionine 0.45%, calcium 1.1%, phosphorus 0.7%, sodium 0,15%, energy 11,4 MJ ME/kg.

Additives per kg:

Nutritional additives: 8,600 i.U. Vitamin A (3a672a), 3.420 i.E. Vitamin D3 (3a671), 45 mg vitamin E (3a700), 12 mg copper as copper (II) sulphate, pentahydrate (E4), 57 mg iron as

iron carbonate (3b101), 60 mg zinc as zinc sulphate, monohydrate (3b605), 80 mg manganese as manganese (II) oxide (3b502), 1.2 mg iodine as calcium iodate, anhydrous (3b202), 0.35 mg selenium as sodium selenite (E8), 2.260 mg hydroxy analogue of methionine (min. 88% total acid).

Technological additives: 17.0 mg butylated hydroxytoluene (BHT) (E321), 7.0 mg Propyl gallate (E310), citric acid (E330), formic acid (E236), lactic acid (E270), propionic acid (E280).

Coccidoistatic and histomonostatic agents: 100 mg monensin sodium (51701).

**Poor diet:**

Ingredients: Crude protein 14.5%, crude fat 4.0%, crude fibre 5.0%, crude ash 6.0%, lysine 0.58%, methionine 0.30%, calcium 1.0%, phosphorus 0.6%, sodium 0,15%, energy 11,4 MJ ME/kg.

Additives per kg:

Nutritional additives: 8,000 i.U. Vitamin A (3a672a), 2.400 i.E. Vitamin D3 (3a671), 16 mg vitamin E (3a700), 12 mg copper as copper (II) sulphate, pentahydrate (E4), 57 mg iron as

iron carbonate (3b101), 60 mg zinc as zinc sulphate, monohydrate (3b605), 80 mg manganese as manganese (II) oxide (3b502), 1.2 mg iodine as calcium iodate, anhydrous (3b202), 0.35 mg selenium as sodium selenite (E8), 1,100 mg hydroxy analogue of methionine (min. 88% total acid).

Technological additives: 17.0 mg butylated hydroxytoluene (BHT) (E321), 7.0 mg Propyl gallate (E310), citric acid (E330), formic acid (E236), lactic acid (E270), propionic acid (E280).

Coccidoistatic and histomonostatic agents: 100 mg monensin sodium (51701).

**Description of the analysis for effects on survival from hatching to day 84**

### We fitted a univariate phenotypic model with ‘survival until the end of the experiment (day 84)’ as response variable, assuming a binary error distribution with a logit link function and fixed the residual variance to one. We modelled random intercepts for pair, mother and father identity. As fixed effects, we fitted diet (categorical variable with two levels: standard and poor), year (categorical variable with two levels: 2019 and 2020), inbreeding coefficient (continuous variable), mother line (categorical variable with two levels: high and low), father line (categorical variable with two levels: high and low), mother replicate (categorical variable with two levels: 1 and 2), father replicate (categorical variable with two levels: 1 and 2), and the 3-way interaction between mother and father line and diet. We did not fit sex as a fixed effect because we could not sex all the individual offspring that died before the end of the experiment.

We did not observe clear differences in survival linked to the diet treatment, although we found a significant effect of inbreeding (Table S1). Individuals with lower inbreeding coefficients were more likely to survive until the end of the experiment than individuals with higher inbreeding coefficients (Table S1). There was no clear effect of year of experiment, or mother and father replicate on survival. The interactive effects of mother and father line with diet were also not important (Table S1).

**Table S1.** Estimates from a univariate phenotypic model on survival from hatching to the end of the study (day 84). Estimates represent posterior modes with associated 95% Credible Intervals (95% CI).

| **Fixed effects** | **Estimate** | **95% CI** |
| --- | --- | --- |
| Intercept | 5.282 | [3.372 , 7.734] |
| Year [2020] | -0.616 | [-1.602 , 0.695] |
| Inbreeding coefficient | -19.27 | [-40.362 , -0.81] |
| Maternal line [low] | -1.419 | [-3.700 , 1.488] |
| Paternal line [low] | -1.618 | [-3.681 , 1.214] |
| Maternal line [low] x Paternal line [low] | 2.885 | [-2.029 , 6.967] |
| Diet [standard] | -0.422 | [-1.614 , 0.877] |
| Diet [standard] x maternal line [low] | 0.994 | [-1.240 , 3.692] |
| Diet [standard] x paternal line [low] | 0.790 | [-1.128 , 2.733] |
| Diet [standard] x paternal line [low] x maternal line [low] | -0.450 | [-3.681 , 3.335] |
| Mother replicate [2] | 0.458 | [-3.200 , 3.576] |
| Father replicate [2] | 0.588 | [-3.369 , 3.189] |
| **Random effects** | **Estimate** | **95% CI** |
| Mother ID | 0.011 | [0.000 , 1.338] |
| Father ID | 0.006 | [0.000 , 1.403] |
| Pair ID | 0.019 | [0.000 , 2.132] |
